# Gap junction communication regulates luminal-myoepithelial crosstalk and cell differentiation in a bilayered human mammary epithelial cell model

**DOI:** 10.1101/2025.09.10.675435

**Authors:** Alec McDermott, Melany Juarez, David Tovar-Parra, Capucine Duval, Shunmoogum (Kessen) Patten, Isabelle Plante

## Abstract

Interactions between luminal and myoepithelial cells are essential for proper mammary gland development and function. However, only a few *in vitro* models allow the study of these interactions and their modulation by extrinsic factors. We developed a layered co-culture system (LCS) that mimics the bilayered architecture of the human breast epithelium, enabling direct contact and bidirectional crosstalk between luminal (MCF-12A) and myoepithelial (MYO1089) cells. We confirmed the formation of adherens and functional gap junctions across the layers. Transcriptomic analysis revealed that co-culture altered gene expression in pathways related to extracellular matrix remodeling, mRNA processing, response to external stimuli, hormonal signaling, receptor activity, and cell cycle regulation. In the LCS, MCF-12A cells exhibited a more luminal-like phenotype, with increased keratin-18 and decreased keratin-14, α-smooth muscle actin (α-SMA), and caldesmon-1 expression compared to 2D monoculture. Conversely, MYO1089 cells showed enhanced expression of myoepithelial markers, including α-SMA, keratin-14, and caldesmon-1. Inhibition of gap junction communication by carbenoxolone disrupted these lineage-specific differentiation patterns. These findings highlight the importance of direct communication in regulating epithelial identity and underscore the LCS as a physiologically relevant model for studying mammary gland biology and external influences that can impact the epithelial differentiation.

**Highlights:** A novel bilayered human co-culture system mimics the luminal and myoepithelial architecture of the mammary epithelium

Direct luminal-myoepithelial contact enables formation of functional adherens and gap junctions

Direct communication via gap junctions is essential for lineage-specific epithelial differentiation

Co-culture induces transcriptional shifts related to proliferation, ECM regulation, and hormone response

## Introduction

The mammary gland comprises two distinct compartments: the stroma and the epithelium (Hennighausen and Robinson, 2005). The epithelium is composed of acini and ducts characterized by a central lumen formed by polarized luminal epithelial cells, surrounded by an outer layer of basal cells, primarily myoepithelial cells (Gudjonsson et al., 2005). Myoepithelial cells were historically perceived as having a limited role in mammary gland development and function, driving only the contraction of the mammary gland during lactation to facilitate milk ejection (Watson and Khaled, 2020). However, it is now well accepted that the myoepithelial layer serves as a mediator for interactions between luminal cells and the stromal components, thereby exerting a profound influence on organ homeostasis and structural integrity (Lanigan et al., 2007) Their primary roles lie in maintaining the precise polarization of luminal cells and participating in tissue remodeling as they secrete components of the basement membrane, such as collagen-IV, laminin-I and -5 and fibronectin, and produce matrix metalloproteinases (MMPs) and their inhibitors (TIMPs) (Gudjonsson et al., 2002, Faraldo et al., 2005, Jolicoeur, 2005, Adriance et al., 2005, Rudolph-Owen and Matrisian, 1998, Barsky and Karlin, 2005, Barsky, 2003). They thus facilitate paracrine regulation and bidirectional communication between the two layers of the epithelium, while simultaneously participating in the intricate process of tissue remodeling (Dickson and Warburton, 1992, Warburton et al., 1982, Rudolph-Owen and Matrisian, 1998). In virgin mice, the functional loss of myoepithelial cells led to extensive branching and the formation of acini-like structures, closely resembling those found in a lactating mammary gland (Moumen et al., 2011). This observation underscores that perturbations in the gene expression of myoepithelial cells have a direct and consequential impact on the proper development of luminal cells within the breast epithelium (Deugnier et al., 2002).

Myoepithelial cells have also been well-characterized as tumor suppressors. Myoepithelial cells form both a physical and chemical barrier, playing a role as defenders against cancer invasion (Adriance et al., 2005, Shams, 2022) by inhibiting the tumorigenic properties of cancer cells, which primarily originate from luminal or stem/progenitor cells (Ingthorsson et al., 2015, Pandey et al., 2010, Sopel, 2010, Polyak and Hu, 2005, Taurin et al., 2006, Skibinski and Kuperwasser, 2015). Consistently, both *in vitro* and *in vivo* studies have shown that early stages of cancer invasion are closely linked to the disruption of myoepithelial integrity (Schnitt, 2009, Winer et al., 2018, Mitchell et al., 2020, Man, 2007). When the myoepithelial layer is compromised, it triggers the secretion of various molecules, including matrix metalloproteinases, tenascin, CXCL12, and CXCL14 (Shams, 2022). These secreted factors can remodel the tumor microenvironment, facilitating interactions between luminal cells and the stroma (Man, 2007, Schnitt, 2009). Additionally, the ability of myoepithelial cells to maintain luminal cell polarity adds another layer of protection against malignancy (Faraldo et al., 2005, Yu et al., 1997). Interestingly, myoepithelial cells near breast carcinomas differ phenotypically from those in normal tissues and may even adopt tumor-promoting characteristics (Barsky and Karlin, 2005, Shams, 2022, Ding et al., 2019), suggesting a dynamic, bidirectional crosstalk between luminal/cancer and myoepithelial cells that influences tumor suppression capacities. Supporting this, disruption of desmosomal adhesion molecules or adherens junctions proteins like E-cadherin leads to improper positioning of luminal and myoepithelial cells in primary culture (Runswick et al., 2001), while inhibition of P-cadherin reduces the tumor-suppressive capacity of myoepithelial cells (Sirka et al., 2018). These findings suggest that bidirectional communication between luminal and myoepithelial cells is essential for epithelial differentiation, and its loss may contribute to breast cancer progression.

Direct cytoplasmic exchange between luminal and myoepithelial cells is also critical for mammary gland development (Biswas et al., 2022). Gap junctions facilitate this exchange, allowing molecules smaller than 1 kDa (e.g., amino acids, nucleotides, glucose, microRNAs, Ca2+, ATP, cAMP, cGMP, IP3) to pass between neighboring cells (Aasen et al., 2016). Gap junctions play roles in proliferation, differentiation, apoptosis, migration, and invasion (Dbouk et al., 2009). Re-expression of Connexin 43 (Cx43) in tumor cells induces redifferentiation, increases epithelial marker cytokeratin 18, decreases mesenchymal marker vimentin, and restores acinar formation in 3D cell cultures (McLachlan et al., 2006). Cx43 re-expression also promotes a less invasive, less tumorigenic phenotype (McLachlan et al., 2007, Hirschi et al., 1996, Qin et al., 2002, Krutovskikh et al., 2000), supporting the idea that Cx43-mediated cytoplasmic exchange promotes proper cell differentiation. Finally, our previous study demonstrated that loss of Cx43 delayed the development of mammary gland, favors hyperplasic mammary gland, and enhances metastasis to the lungs in a loss-of-function mice model (Plante et al., 2008; Plante et al., 2011), further supporting a crucial role of Cx43 in mammary gland biology.

In line with the Tox21 initiative to reduce animal testing and improve chemical safety evaluation using alternative methods, there is a pressing need for physiologically relevant *in vitro* models. Understanding the crosstalk between luminal and myoepithelial cells is key to evaluating how environmental exposures influence mammary development, and how dysregulation of this crosstalk relates to breast cancer. We hypothesize that bidirectional communication between luminal and myoepithelial cells is necessary for proper epithelial differentiation. Here, we present a robust layered co-culture system (LCS) that models the human mammary epithelium and allows investigation of cell-to-cell direct interactions critical to epithelial phenotype. This model provides a valuable platform for mechanistic and toxicological studies.

## MATERIALS AND METHODS

### Cell culture

MCF-12A cells (ATCC CRL-10782TM) were purchased from ATCC (Manassas, VA, USA). These are non-tumorigenic human mammary epithelial cells derived from adherent cells in a mixed population of breast tissue. MYO1089 cells were generated by Dr. Mike O’Hare (University of London, UK) (O’Hare et al., 1991) and kindly donated by Dr. Louise J. Jones (Barts Cancer Institute, Queen Mary, University of London). They were derived from a human breast reduction mammoplasty tissue and immortalized via genomic incorporation of the SV40 large T antigen gene. MCF-12A cells were maintained in phenol red-free Dulbecco’s Modified Eagle’s Medium/Ham’s F12 (DMEM/F12; 21041025, Thermo Fisher Scientific) supplemented with 5% (v/v) horse serum (16050-122, Thermo Fisher Scientific), 20 ng/mL human recombinant epidermal growth factor (hEGF; E9644, Sigma-Aldrich), 10 µg/mL bovine insulin (I6634, Sigma-Aldrich), and 500 ng/mL hydrocortisone (7904, STEMCELL Technologies), and propagated according to ATCC guidelines. MYO1089 cells were maintained in Ham’s F12 medium (11765054, Thermo Fisher Scientific) supplemented with 10% fetal bovine serum (FBS; 12484-028, Thermo Fisher Scientific), 1 µg/mL hydrocortisone (7904, STEMCELL Technologies), 5 µg/mL bovine insulin (I6634, Sigma-Aldrich), and 10 ng/mL hEGF (E9644, Sigma-Aldrich), following the same propagation protocol as MCF-12A cells. Cells were incubated at 37°C with 5% CO₂. For all experiments, both MCF-12A and MYO1089 cells were cultured in MCF-12A medium to reduce media-related bias. This media change for MYO1089 did not affect their morphology, expression of specific markers or doubling time.

### Layered Co-Culture System (LCS) Setup

High pore density 3.0 μm pore size PET (polyethylene terephthalate) track-etched membrane cell culture inserts (353492/353092, Falcon, Corning) were used in the development of this model. This pore size was intentionally selected to enable direct cellular interactions between the two cell layers while preventing cell migration across the membrane. Immunofluorescence analyses were performed using 24-well-plate size inserts, while for other analyses, 6-well-plate (P6) size inserts were preferred. MYO1089 cells were harvested using 0.25% trypsin-EDTA (252000-072, Thermo Fisher Scientific) and seeded on the underside of inverted inserts at a density of 500,000 cells in 500 μL of media for 6-well inserts or 300,000 cells in 200 μL for 24-well inserts (Figure 1A, Supplementary Figure 1A-B). Inserts were maintained at 37°C and 5% CO₂ for 6 hours to allow cell adhesion. To maintain sterility during this adhesion period, the inserts were assembled in an inverted 6-or 24-well plate configuration (Supplementary Figure 1A-B). Specifically, inverted inserts were placed on the lid of a corresponding well plate, and the base of the plate was used as a cover, preserving media through surface tension and preventing desiccation (Supplementary Figure 1C). After 6 hours, the inserts and plates were returned to their upright position, and MCF-12A cells were seeded on the upper surface at the same density as the MYO1089 cells. MCF-12A culture medium was added to ensure full coverage of both compartments. The co-culture system was incubated at 37°C and 5% CO₂ for 16–18 hours before further analysis.

**Figure 1.**
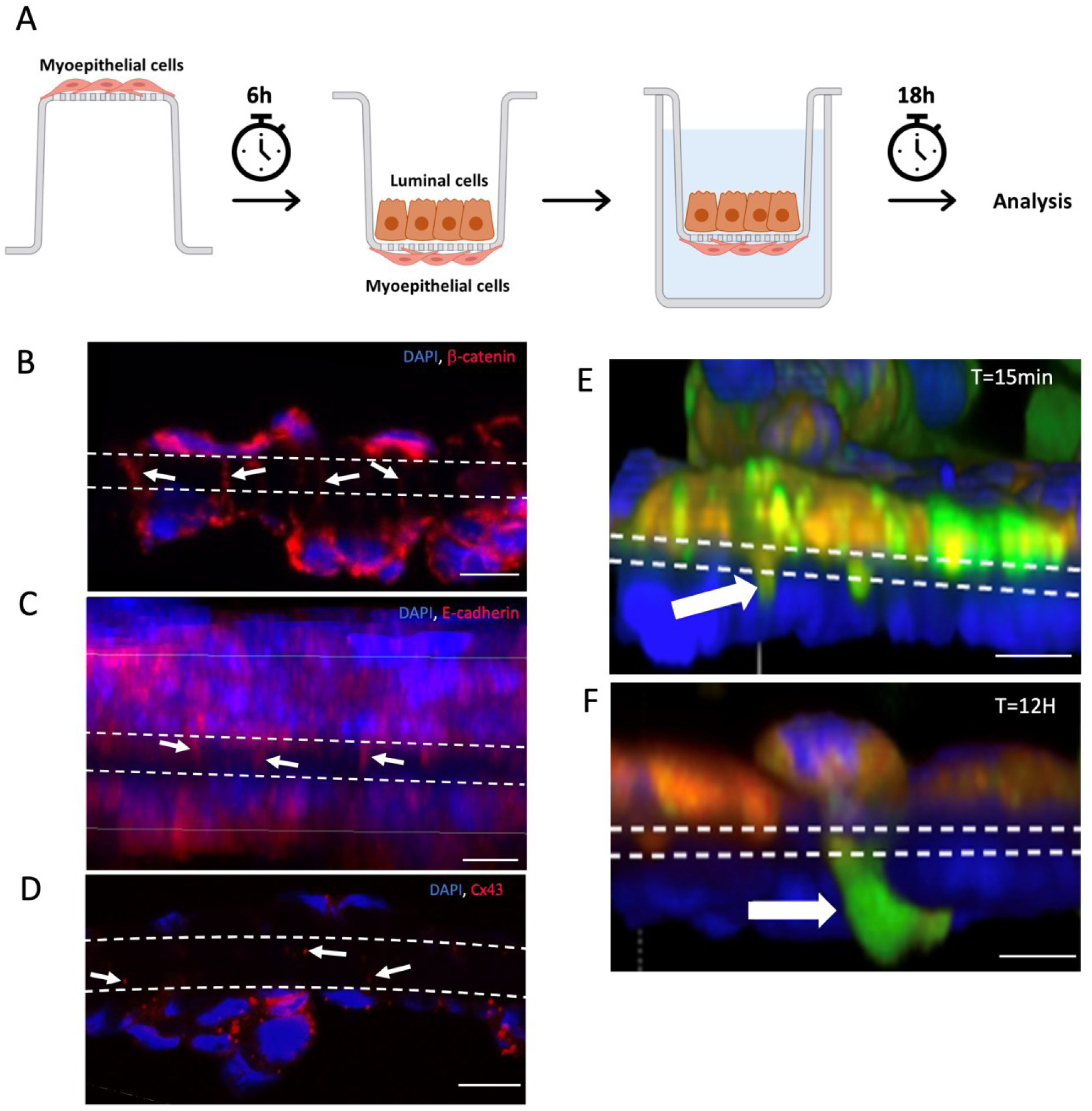
Cells from each layer are able to form direct contact and communicate via the pores of the inserts. (A) General workflow of the Layered-Culture System (LCS). MYO1089 cells are seeded on the bottom side of the membrane of an inverted insert and allow to adhere for 6 hours. After the insert is flipped and MCF-12A cells are seeded in the top side of the membrane, and placed in a plate with media. For cytometer settings and GJIC assays, the MCF-12A cells are preloaded with dyes, and MCF-12A cells are added to the bottom of the plate to evaluate indirect communication. Cells are maintained in co-culture for approximately 16 to 18 hours prior to analyses. (B) 8 μM cross-section showing β-catenin projection and direct contacts between MCF-12A and MYO1089. (C) Side view of a 3D representation of a Z-stack showing E-cadherin through the pores of the membrane. (D) 8 μM cross-section showing the presence of Cx43 within the pores of the membrane (white arrows). (E) Calcein (green) and DiL (Red) dyes were loaded in donor cells (MCF12-A), and calcein, but not DiL, could be observed in MYO1089 cells after 15 minutes and 12 hours of co-culture (E, F). For all images, Nuclei are in blue, Scale bar = 10 μm. MCF-12A luminal cells are seeded on top of the insert and MYO1089 cells on the other side and the white dashed lines delimit the position of the porous membrane.

### Gap Junction Intercellular Communication Assays

For dye transfer analysis, we used two dyes as described in previous studies (Goldberg et al., 2002): calcein AM (554217, Corning) and 1,1ʹ dioctadecyl 3,3,3ʹ,3ʹ tetramethylindodicarbocyanine perchlorate (DiL; 08774502, Corning) (Goldberg et al., 2002). Calcein-AM is a membrane permeable-green fluorescent dye that, once internalized, is hydrolyzed by intracellular esterases into fluorescent calcein and becomes trapped inside the cell. It can then pass to adjacent cells only via functional gap junctions (Mariappan et al., 1999). In contrast, DiL is a lipophilic red fluorescent dye that stains cell membranes and does not transfer between cells, serving as a negative control for gap junction communication (Warawdekar, 2019). MCF-12A cells were used as donor cells and labelled by incubation in medium containing both 5 µM calcein-AM and 0.072 µM DiL for 30 minutes at 37°C and 5% CO₂. Dye concentrations were optimized to avoid cytotoxicity while ensuring quantifiable signal in both donor and recipient cells. After incubation, cells were washed with PBS, trypsinized, centrifuged (125 × g, 7 minutes), and washed three times in PBS. The labelled MCF-12A cells were then resuspended in culture medium, counted, and seeded on the upper side of porous membrane inserts that had been pre-seeded on the underside with unstained MYO1089 cells. Co-cultures were maintained for 16–18 hours before analysis by either flow cytometry or confocal microscopy. To confirm that dye transfer was mediated by gap junctions and not by diffusion through the media, an additional control layer of MCF-12A cells was seeded on the well bottom (Figure 1A).

### Gap Junctional Intercellular Communication Inhibition

Carbenoxolone disodium salt (CBX; AAJ6371403, Thermo Scientific) was used to inhibit gap junction intercellular communication (GJIC). A 20 000 µM stock solution of CBX was prepared in PBS, which also served as the vehicle control. Cytotoxicity of CBX was first assessed by treating cells with concentrations ranging from 0 to 100 µM for 24 hours. For inhibition of GJIC during dye transfer experiments, donor MCF-12A cells were labelled with calcein-AM as described above. After labelling, the cells were washed three times with PBS and incubated with 100 µM CBX for 6 hours. In parallel, MYO1089 receiving cells were seeded on the underside of the insert and also treated with 100 µM CBX. After the pre-incubation period, the inserts were flipped to the upright position and labelled donor cells were seeded onto the upper membrane surface. Co-cultures were maintained for 16–18 hours in the presence of either 100 µM CBX or the PBS vehicle control prior to analysis.

### Flow cytometry

After 16–18 hours of incubation, inserts containing calcein-labelled MCF-12A (donor) cells and MYO1089 (receiving) cells on opposite sides of the membrane were washed with PBS. Cells on each side were sequentially collected using 0.25% trypsin and a cell scraper, carefully separating cells from the upper and lower surfaces of the membrane. Cells were diluted in culture medium, centrifuged at 125 × g for 7 minutes, and resuspended in 0.5 mL of PBS. Flow cytometric analysis was performed using a NovoCyte Penteon cytometer (Agilent), acquiring 50 000 singlet events per sample. As an internal control, MCF-12A cells seeded at the bottom of the well (not in direct contact with MYO1089 cells) were recovered and analyzed to verify that dye transfer occurred via direct gap junctional communication and not through leakage into the medium (Figure 1A). Data analysis was performed using FlowJo VX software.

### Immunofluorescence of cryosections

Membranes from 6-well (P6) inserts were carefully removed using a scalpel and embedded in Scigen Tissue-Plus™ O.C.T. Compound (23-730-571, Fisher Scientific). Transversal 8 µm cryosections were prepared using a cryostat (Microm HM525, Thermo Scientific) and fixed in a solution of 80% methanol and 20% acetone for 10 minutes, followed by two PBS washes. Sections were blocked with 2% BSA and 0.1% Triton X-100 in PBS for 60 minutes at room temperature, then incubated for 2 hours with primary antibodies diluted in blocking solution (see Supplementary Table 1). Following three 5-minute PBS washes, sections were incubated with respective secondary antibodies for 1 hour at room temperature (Supplementary Table 1). Nuclei were stained with DAPI (1 µg/mL) for 5 minutes, then sections were mounted in Fluoromount-G (SouthernBiotech, 0100-01). Confocal images were captured using a Nikon A1R+ confocal microscope and processed with NIS-elements software.

### Confocal Imaging of Whole Membranes and 2D Cell Cultures

Immunofluorescence analysis was performed on whole membranes following co-culture using the LCS as previously described. After 16 hours, the cell culture medium was removed, and membranes were washed with PBS. Cells were then fixed with 4% formaldehyde for 15 minutes, followed by two washes with PBS. Blocking and antibody incubations were conducted as detailed previously (Supplementary Table 1). After incubation with secondary antibodies, membranes were washed three times for five minutes each. Nuclear staining was performed using DAPI (1 µg/mL) for 5 minutes at room temperature, followed by a final PBS wash. Subsequently, membranes were carefully excised with a clean scalpel and mounted between two 22 mm × 30 mm coverslips using Fluoromount-G (SouthernBiotech). Whole membranes were also used to assess DiL and calcein transfer. Cells were labelled with the respective dyes, seeded as described, and fixed following the same protocol. Membranes were excised and mounted according to the method outlined above.

For 2D cell cultures, cells were seeded on 35 mm glass-bottom dishes (P35GC-0-14-C, MatTek) and cultured until confluence. Immunollabeling was then performed as described above. Finally, Fluoromount-G was applied to fix the coverslip onto the cell layer.

### RNA Extraction and Sequencing

After co-culture, cells from each side of the membrane were harvested separately for RNA extraction. First, the culture medium was removed, and inserts were washed with PBS. Cells were detached as previously described, and fresh media was used to independently recover cells from each side of the membrane. The recovered cells were centrifuged at 125 × g for 7 minutes. In parallel, MCF-12A and MYO1089 cells from monoculture (single cell type) were collected as controls to assess the effects of co-culture. Cell pellets were processed for total RNA extraction using the Aurum™ Total RNA Mini Kit (7326820, Bio-Rad) according to the manufacturer’s instructions. RNA quantity and quality were assessed prior to sequencing using the RNA 6000 Nano kit (5067–1511, Agilent) and the 2100 Bioanalyzer system (G2939BA, Agilent). Samples with an RNA Integrity Number (RIN) ≥ 8 were submitted to the Montreal Clinical Research Institute (IRCM) sequencing facility. Ribo-depleted libraries were prepared and sequenced on an Illumina NovaSeq 6000 system with a sequencing depth of 50 million reads per sample. Raw sequencing read quality was evaluated using FASTQC v0.11.8. No trimming was necessary based on quality assessment. Reads were aligned to the human reference genome (GRCh38) using STAR v2.7.6a, achieving an average of 90% uniquely mapped reads. Raw gene counts were generated using FeatureCounts v1.6.0, referencing the Ensembl genome database (release 108). Differential gene expression analysis was performed separately for each cell type using the DESeq2 R package (v1.42.1), controlling for batch effects related to RNA extraction date. P-values were adjusted for multiple testing using the Benjamini-Hochberg method. Heatmaps of differentially expressed genes (DEGs) were generated based on the z-scores of normalized counts after batch effect correction with Limma’s removeBatchEffect function. Functional enrichment analysis of DEGs, including Gene Ontology and pathway analysis, was conducted with the gprofiler2 R package (Raudvere et al., 2019). All bioinformatics analyses were carried out at the Bioinformatics Core Facility of the Montreal Clinical Research Institute (IRCM). Sequencing data have been deposited in NCBI under accession number PRJNA1268018.

### RT-qPCR

Following exposure to CBX or vehicle control, MCF-12A and MYO1089 cells in the LCS were recovered, and RNA was extracted as described previously. After validating RNA quality, complementary DNA (cDNA) was synthesized using the iScript cDNA Synthesis Kit (1708891, Bio-Rad) according to the manufacturer’s instructions. Quantitative PCR analyses were performed using the CFX96 Touch Real-Time PCR Detection System (Bio-Rad). Primers were designed with Primer-BLAST (https://www.ncbi.nlm.nih.gov/tools/primer-blast/, last accessed May 13, 2025) and their quality was validated using OligoAnalyzer (https://www.idtdna.com/calc/analyzer, Integrated DNA Technologies, last accessed May 13, 2025). Primers’ sequences are provided in Supplementary Table 2. qPCR assays were conducted using SsoAdvanced SYBR Green Supermix (1725274, Bio-Rad). Data analysis was performed with the CFX Maestro software. Primer efficiency and specificity were assessed for each primer pair using standard curve analysis and dissociation curve (melting curve) analysis of PCR products. Reference genes were selected based on the geNorm algorithm. Relative gene expression levels were quantified using the comparative cycle threshold (ΔΔCt) method.

### Western Blot

Protein extraction was performed from fresh cells cultured in P6 wells, which had been exposed to either vehicle control (PBS diluted 1:2000 in culture medium) or 100 µM CBX for approximately 18 hours. Cells were harvested using trypsin as described previously. The resulting cell pellet was resuspended in cold triple-detergent lysis buffer (50 mM Tris, 150 mM NaCl, 0.02% sodium azide, 0.1% SDS, 1% Nonidet P40, 0.5% deoxycholate, pH 8), supplemented with 1.25 M sodium fluoride, 1 M sodium orthovanadate, and a protease/phosphatase inhibitor cocktail (Halt Protease and Phosphatase Cocktail; 78447, Fisher Scientific). Mechanical disruption was achieved by repeated pipetting. Samples were centrifuged at 13,000 × g for 10 minutes at 4°C, and the supernatant was collected, aliquoted, and stored at −80°C until analysis. Protein concentration was determined using the Pierce™ BCA Protein Assay Kit (23225, Thermo Fisher Scientific). For semi-quantitative Western blotting, equal amounts of total protein were loaded onto 10% TGX Stain-Free™ acrylamide gels (1610183, Bio-Rad). Following electrophoresis, proteins were transferred to PVDF membranes using the Trans-Blot Transfer System (Bio-Rad). Membranes were imaged with the ChemiDoc MP Imaging System (Bio-Rad) for total protein normalization. Membranes were blocked with 0.1% TBS-Tween containing 5% dry milk, followed by overnight incubation at 4°C with Connexin 43 primary antibody (C6219, Sigma Aldrich) diluted 1:750. After washing with 0.1% TBS-Tween, membranes were incubated with anti-rabbit IgG HRP-linked secondary antibody (7074, Cell Signaling) at a 1:10,000 dilution. Signal detection was performed using Clarity™ Western ECL Blotting Substrate (1705061, Bio-Rad), and chemiluminescent signals were captured using the ChemiDoc MP Imaging System. Band intensities were normalized to total protein levels in the corresponding lanes.

### Statistical analysis

Measurement values were expressed as a mean and standard deviation (±SD). Statistical analyses were conducted using ANOVA and T-tests when parametric assumptions were met. If these assumptions were not met, the Kruskal-Wallis test and Mann-Witney Test were used. Statistically significant differences were represented using the following p-values *p<0.05, **p<0.01, ***p<0.001, ****p<0.0001, “ns” indicated the absence of a significant difference. GraphPad Prism software version 10.0 was used for statistical analysis and graphical representation. Experiments were conducted in triplicate and independently replicated at least three times.

## RESULTS

### Luminal and Myoepithelial Cells form Physical and Functional Interactions Across a Porous Membrane

Our objective was to establish a model to study the bidirectional crosstalk between luminal and myoepithelial cells within the mammary gland epithelium, with applications in development, cancer progression, and toxicology. To this end, we co-cultured luminal MCF-12A cells and myoepithelial MYO1089 cells on opposite sides of a porous membrane containing 3 μm pores (Figure 1A).

We first assessed whether physical interactions between the two cell types occurred. Immunofluorescence staining of membrane cross-sections revealed cell-cell contact both within each layer and through the porous membrane (Figure 1B, C, D). Specifically, β-catenin and E-cadherin were detected at the plasma membranes of both luminal and myoepithelial cells, as well as within the insert’s membrane pores, indicating the potential formation of adherens junctions likely mediated by cell projections spanning the insert’s membrane (Figure 1B, C). Furthermore, the presence of Cx43 within the membrane pores was confirmed (Figure 1D), suggesting that cytoplasmic exchange of small molecules via gap junctions likely occur.

To evaluate functional communication through gap junctions, we adapted a well-characterized dye transfer assay for use in the LCS (Warawdekar, 2019). Luminal cells were labelled with calcein-AM (a gap junction-permeable dye) and DiL (a membrane-bound dye), then co-cultured with myoepithelial cells for up to 12 hours. After fixation and nuclear staining with DAPI, calcein fluorescence was observed within the membrane pores as early as 15 minutes post co-culture (Figure 1E). After 12 hours, calcein-positive cells were detected in the bottom (myoepithelial) layer, confirming effective GJIC (Figure 1F). Importantly, DiL labelling confirmed that donor luminal cells remained confined to their original side of the membrane, indicating that no cell migration occurred across the pores (Figure 1F).

### RNA-seq Reveals Distinct Transcriptomic Profiles of MCF-12A and MYO1089 Cells in Layered Co-Culture Compared to 2D Monoculture

Building on the evidence that luminal and myoepithelial cells can form direct communication through the pores of the membrane, we next sought to examine the transcriptomic changes associated with this interaction, as it is well established that interactions between luminal and myoepithelial cells are essential for proper mammary gland development and function (Moumen et al., 2011, Faraldo et al., 2005, Deugnier et al., 2002). It is well established that interactions between luminal and myoepithelial cells are essential for proper mammary gland development and function (Moumen et al., 2011, Faraldo et al., 2005, Deugnier et al., 2002). We showed that luminal and myoepithelial cells can form direct communication through the pores of the membrane, we thus next sought to examine the molecular signatures associated with this interaction. Using RNA sequencing, we performed a transcriptomic analysis between harvested cells from 2D monoculture and LCS.

The principal component analysis (PCA) plots (Figure 2A and Figure 3A) clearly illustrate differences in gene expression profiles between cells grown in co-culture (cc) and those maintained in 2D monoculture. Importantly, all cells were cultured in the same media conditions to avoid potential confounding effects from differences in culture medium composition.

**Figure 2.**
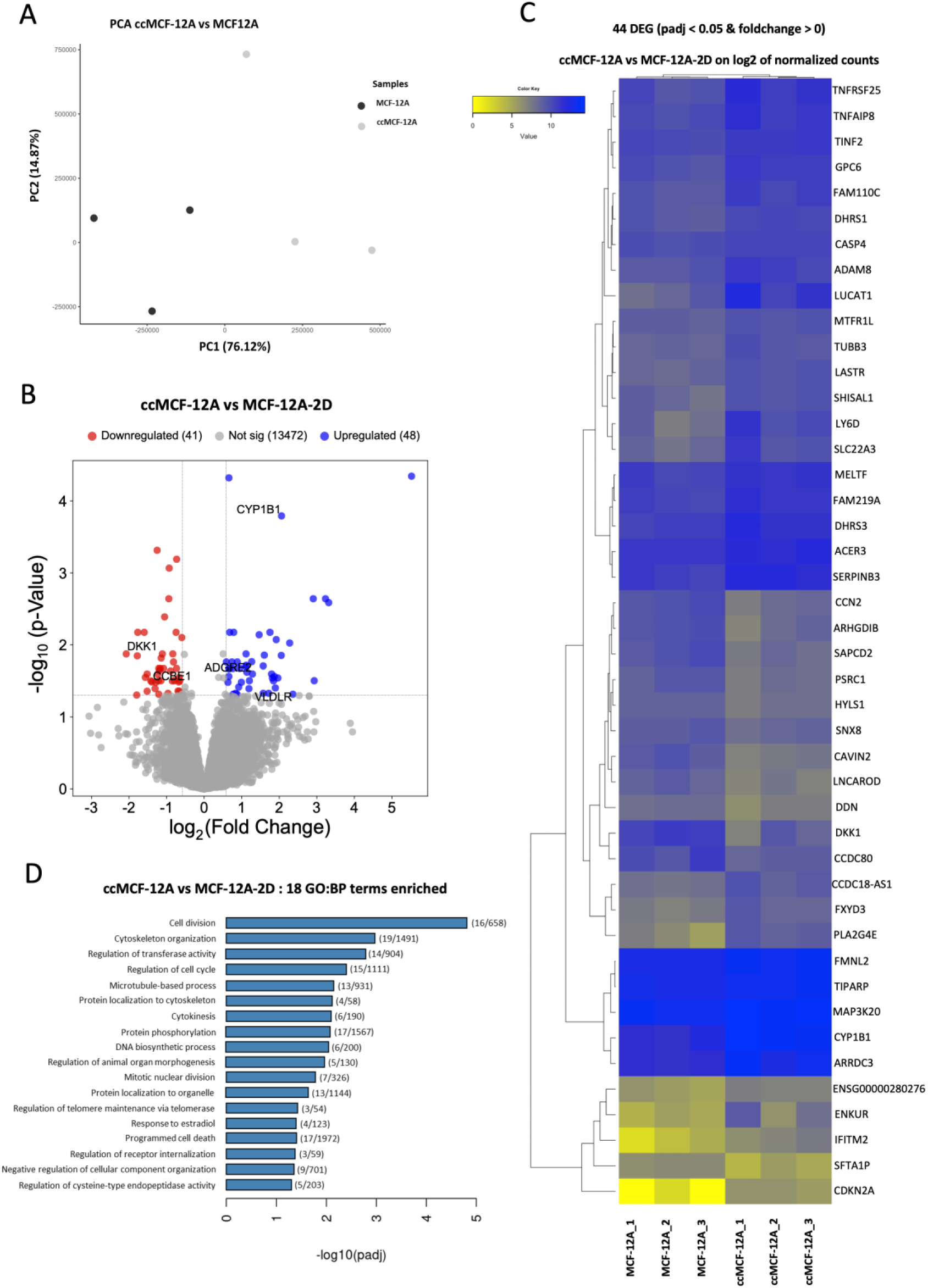
**Total RNA sequencing of co-cultured MCF-12A compared to classical 2D-cultured MCF-12A suggests a more differentiated phenotype**. (A) Principal Component Analysis (PCA) of normalized RNA-seq read counts of co-cultured MCF-12A (ccMCF-12A) vs MCF-12A cultured in a classical way (MCF-12A). (B) Volcano Plot of DESeq2 analysis. Data was considered significant when adjusted q-value ≤ 0.05 and fold change ≥ 1.5. (C) Heatmap representation of significantly different gene expression of log2 of normalized counts (padj ≤ 0.05) and fold change different than 0. (D) Deregulated Biological Process between conditions. (# of deregulated genes / # of genes within the pathway).

**Figure 3.**
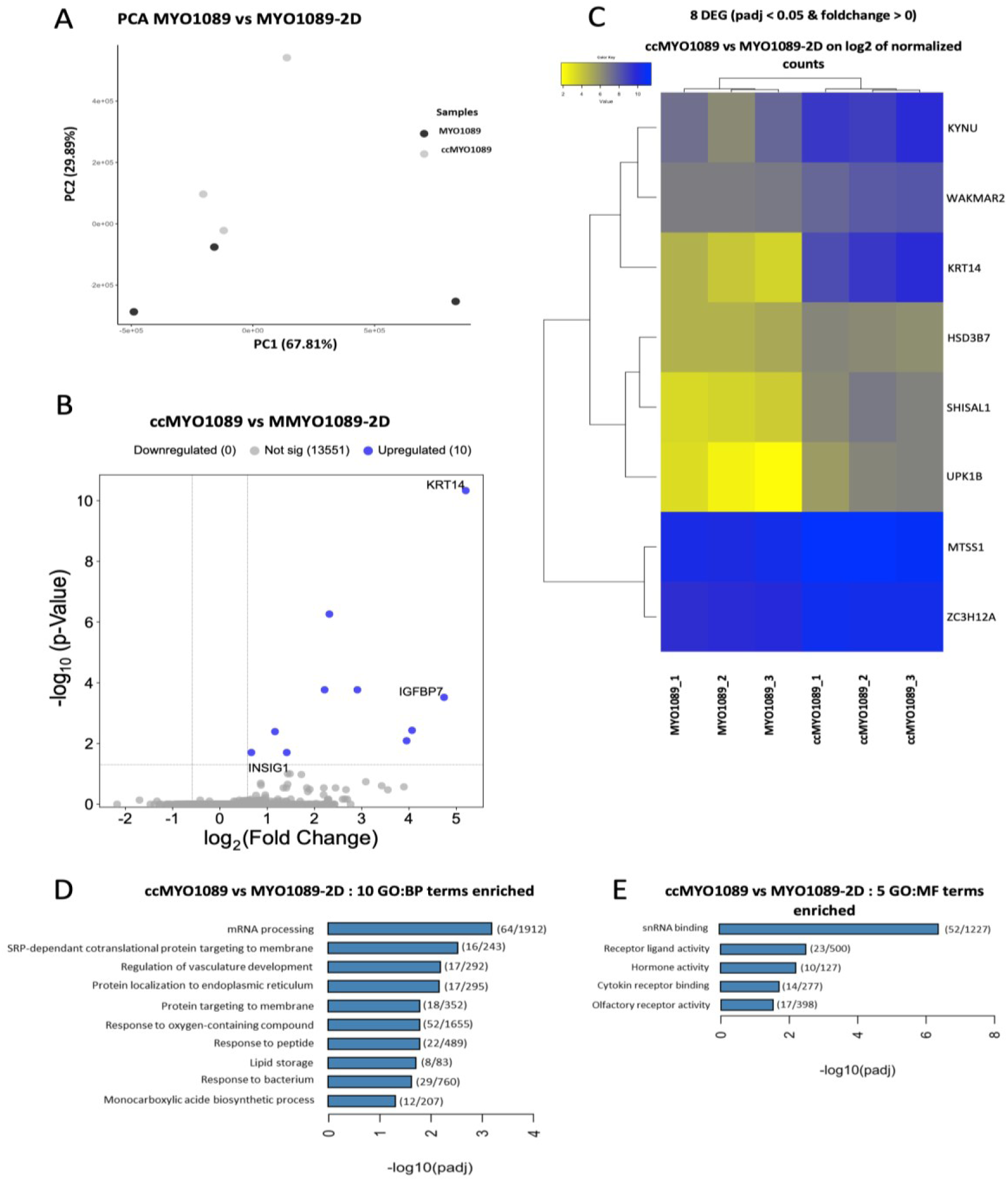
Total RNA sequencing of co-cultured MYO1089 compared to classical 2D-cultured MYO1089 suggests a more differentiated phenotype. (A) Principal Component Analysis (PCA) of normalized RNA-seq read counts of co-cultured MYO1089 (ccMYO1089) vs MYO1089 cultured in a classical way (MYO1089). (B) Volcano Plot of DESeq2 analysis. Data was considered significant when adjusted q-value ≤ 0.05 and fold change ≥ 1.5. (C) Heatmap representation of significantly different gene expression of log2 of normalized counts (padj ≤ 0.05) and fold change different than 0. (D) Deregulated Biological Process between conditions. (# of deregulated genes / # of genes within the pathway). (E) Deregulated Molecular Functions between conditions. (# of deregulated genes / # of genes within the pathway).

When comparing co-cultured MCF-12A cells (ccMCF-12A) with their 2D monoculture counterparts (MCF-12A-2D), a total of 20 867 genes showed significant differential expression at a p-value threshold of < 0.05. However, applying a fold-change cutoff greater than 1.5 and filtering for adjusted p-values yielded only 89 genes (48 upregulated and 41 downregulated) with notable changes (Figure 2B). A comparable pattern was observed for the myoepithelial MYO1089 cells, where 473 genes were significantly differentially expressed between ccMYO1089 and MYO1089-2D; yet, only 10 upregulated genes passed the fold-change and adjusted p-value thresholds (Figure 3B).

To gain a deeper insight on these transcriptomic differences, heatmaps were generated using log2-transformed normalized counts filtered for significantly differentially expressed genes (adjusted p-value < 0.05) with fold-changes different from zero. The heatmap for MCF-12A cells (Figure 2C) revealed 44 differentially expressed genes capable of effectively clustering the co-culture and monoculture samples, underscoring a marked shift in gene expression profile induced by co-culture conditions. Conversely, the heatmap comparing ccMYO1089 and MYO1089-2D cells (Figure 3C) identified only 8 genes capable of clustering the two groups, suggesting that the impact of co-culture on the transcriptome of myoepithelial cells is comparatively less pronounced than in luminal cells.

To interpret the biological relevance of these gene expression changes, we performed Gene Ontology (GO) enrichment analyses focusing on both Biological Process (BP) and Molecular Function (MF) categories. In ccMCF-12A cells, 18 enriched BP terms were identified, the majority of which were related to cell division processes and cytoskeleton organization. Interestingly, these cell division-related genes are predominantly downregulated (Figure 2D), consistent with the known inverse relationship between cellular proliferation and differentiation. Within MF, only one significant enrichment was observed: “fibronectin binding,” which plays a critical role in regulating cell-extracellular matrix (ECM) interactions (data not shown).

For ccMYO1089 cells, GO enrichment analysis revealed 10 significant BP terms, predominantly linked to lipid storage, monocarboxylic acid biosynthetic process, and mRNA maturation processes occurring within the cell nucleus and to cellular responses to external stimuli (Figure 3D). The MF analysis in this cell type yielded five enriched terms, with notable emphasis on mRNA maturation, as well as terms associated with hormonal activity and receptor binding (Figure 3E).

To validate the RNA-seq results, we conducted RT-qPCR for upregulated genes like CYP1B1 (Cytochrome P450 Family 1 Subfamily B Member 1), ADGRE2 (Adhesion G Protein-Coupled Receptor E2), and VLDLR (Very-low-density lipoprotein receptor), and downregulated genes like DKK1 (Dickkopf Wnt Signaling Pathway Inhibitor 1) and CCBE1 (Collagen and Calcium Binding EGF Domains 1) in the luminal comportment of the LCS. The results demonstrated a differential gene expression level in the co-culture MCF12A system compared to MCF12A 2D (Supplementary Figure 2A-B). To further validate these findings, we performed RT-qPCR for KRT14 (Keratin 14), IGFBP7 (Insulin-Like Growth Factor Binding Protein 7), and INSIG1 (Insulin-Induced Gene 1) genes for the myoepithelial cells from the LCS compared to 2D monoculture (Supplementary Figure 2C). The data observed by RT-qPCR follows the same expression pattern found in the RNAseq data.

### MCF-12A Luminal and MYO1089 Myoepithelial Cells Communicate via Gap Junctions through the Porous Membrane to Establish Direct Cell-Cell Communication

To quantitatively assess direct communication through gap junctions in the LCS, luminal MCF-12A cells were preloaded with calcein-AM prior to seeding, then allowed to interact with MYO1089 myoepithelial cells over a 16–18 hour incubation period. Following this, the two cell populations were separately harvested and analyzed by flow cytometry (Figure 4).

**Figure 4.**
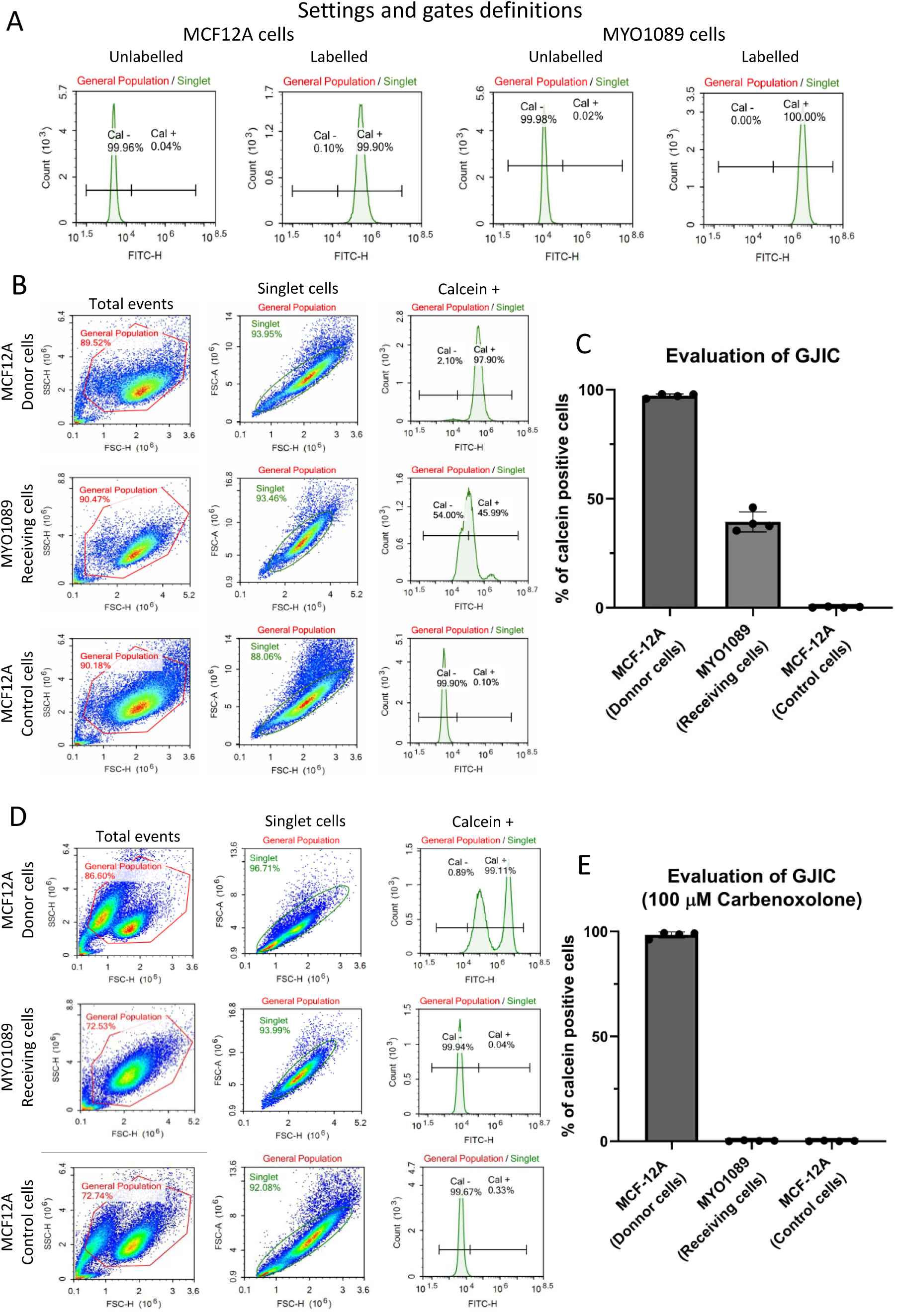
Quantification of gap junction activity between luminal MCF-12A and myoepithelial MYO1089 cells. Luminal cells (donor cells) were pre-stained with calcein-AM and co-cultured in LCS with myoepithelial cell (receiving cells) for 16h to 18h. Luminal and myoepithelial cells were harvested and analyzed through flow cytometry after 16 hours of co-culture. (A) Representation of the gate adjustment for calcein positive and negative MCF-12A and MYO1089 cells. (B) Representative analysis of the communication between MCF-12A and MYO1089. MCF-12A (Donnor cells) are calcein positive cells colored prior to the co-culture. MYO1089 (Receiving cells) are not colored and placed on the bottom side of the insert and allowed to be in direct communication with the colored MCF-12A. MCF-12A (Control cells) are not colored and placed at the bottom of the well, as a control for dye that leaches into the media. (C) Almost all MCF-12A (Donor cells) cells are calcein positive (96 ± 1%), while 39 ± 4% of MYO1089 (Receiving cells) are positive for calcein, demonstrating efficient GJIC between the two layers of cells. MCF-12A (Control cells) cells at the bottom of the well are negative for calcein (0.3 ± 0.3%), confirming that calcein is not transferred through the medium. Graphs represent the mean +/-SD of four independent assays. (D) Representation of the communication between MCF-12A and MYO1089 exposed to a gap junction inhibitor (100 μM of carbenoxolone). (E) Carbenoxolone-exposed MCF-12A (Donor cells) remained calcein positive (98 ± 1.5%%), while 0.25 ± 0.2% of MYO1089 (Receiving cells) are positive for calcein upon exposure to carbenoxolone, confirming the inhibition of GJIC. MCF-12A (Control cells) cells at the bottom of the well (Control cells) are still negative for calcein (0.1 ± 0.1%). Graphs represent the mean +/-SD of four independent assays.

Flow cytometry parameters and gating were carefully established using calcein-labelled and unlabelled cells to minimize bias (Figure 4A). As expected, the donor MCF-12A cells in the upper chamber retained a high proportion of calcein-positive cells (96 ± 1%) (Figure 4B, C). Importantly, approximately 39 ± 4% of the MYO1089 cells in the receiving layer, initially unlabelled, were found to be positive for calcein after co-culture, confirming efficient gap junction intercellular communication (GJIC) between the two cell types across the porous membrane (Figure 4B, C).

To rule out the possibility that calcein-positive MYO1089 cells resulted from dye leakage into the media, MCF-12A cells located at the bottom of the well (separate from the upper-layer co-culture) were also harvested and analyzed. In this control group, only about 0.3 ± 0.3% of MCF-12A cells were calcein-positive, demonstrating that dye transfer to the receiving MYO1089 cells occurs predominantly via direct cytoplasmic exchange through gap junctions rather than extracellular dye diffusion (Figure 4B, C). These findings validate the LCS as a reliable *in vitro* model for studying GJIC and direct cell-cell interactions between luminal and myoepithelial cells of the bilayered mammary epithelium.

To further validate that GJIC was mediating this communication, we evaluated the effect of carbenoxolone (CBX), a known pharmacological inhibitor of gap junction channels. Inserts containing the three cell compartments—MCF-12A donor cells, MYO1089 receiving cells, and MCF-12A control cells at the bottom of the well—were treated with 100 µM CBX. Post-treatment, nearly all donor MCF-12A cells (98 ± 1.5%) were able to uptake and metabolize calcein-AM into fluorescent calcein (Figure 4D, C), confirming that CBX does not inhibit dye uptake or cellular viability. However, CBX treatment dramatically inhibited dye transfer between cell layers, with only 0.25 ± 0.2% of MYO1089 receiving cells staining positive for calcein, indicating effective blockade of GJIC (Figure 4D, E). As expected, MCF-12A control cells at the bottom of the well remained unstained, ruling out indirect dye transfer (Figure 4D, E).

Importantly, while 100 µM CBX effectively inhibits GJIC, it does not appear to be cytotoxic, as the metabolization of calcein-AM requires viable cells. Of note, CBX treatment also induced morphological changes in the cells, evidenced by the appearance of two distinct fluorescent peaks in the flow cytometry analysis of MCF-12A donor cells (Figure 4D). This pattern was corroborated by imaging of 2D cultured cells treated with CBX (Supplementary Figure 3A).

Additionally, Western blot analysis revealed a significant reduction in Cx43 protein levels in both MCF-12A and MYO1089 cells following CBX treatment in 2D culture (Supplementary Figure 3B-C), suggesting that CBX likely inhibits Cx43 function by down-regulating the protein levels (Liu et al., 2022). CBX does not alter the transcriptional gene expression of Cx43 at the mRNA level (Supplementary Figure 3D). This inhibition of GJIC within the cell layer corresponds with altered cell morphology, supporting the concept that proper gap junction communication is essential for maintaining normal cell phenotype and function. Exposure to CBX altered some of the relative gene expression validated by RT-qPCR (Supplementary Figure 2 A-C). This suggests that gap junction plays a critical role in cell differentiation, but direct contact may also be important.

### Direct Cell-Cell Communication Drives Differentiation of MCF-12A and MYO1089 Cells

It is well known that the interaction between luminal and myoepithelial cells is essential for proper mammary gland development and function. In addition, our unbiased transcriptomic data (Figures 2-3) suggested that it also enhanced their differentiation. Therefore, we examined potential alterations in protein expression of specific markers when the two cell populations were maintained in co-culture. We observed that changes in protein expression in luminal and myoepithelial cells were dependent on the culture conditions. When cultured as monocultures under traditional 2D conditions, both MCF-12A luminal and MYO1089 myoepithelial cells appeared poorly differentiated, as they expressed markers characteristic of both lineages. Specifically, both cell lines expressed myoepithelial markers keratin 14 (K14), caldesmon-1, and α-smooth muscle actin (α-SMA), as well as the luminal marker keratin 18 (K18) (Figure 5A, MCF-12A 2D and MYO1089 2D).

**Figure 5.**
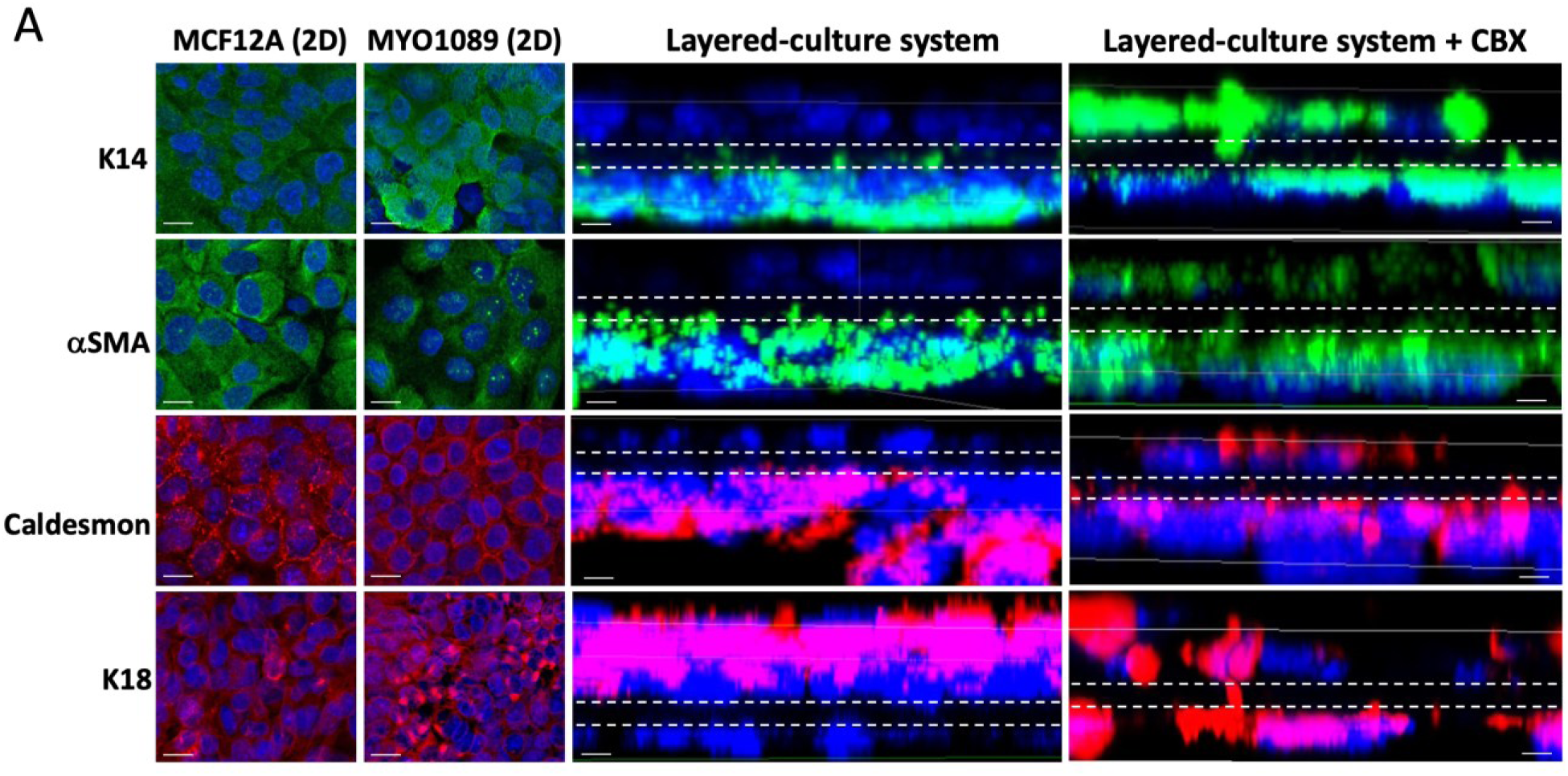
Gap junction-mediated communication between MCF-12A and MYO1089 is required for proper cell differentiation. (A) Representation of immunofluorescence of classical 2D cell culture (MCF-12A 2D and MYO1089 2D), cells co-cultured in the Layered-culture system (LCS) and exposed to 100 μM of carbenoxolone co-cultured in the LCS. Both MCF12-A and MYO1089 cells showed positive staining for K14, α-SMA, caldesmon-1 and K18 when cultured in Classic 2D culture conditions, but gained a more differentiated phenotype when co-cultured in LCS as defined by positive staining for K18 only in MCFR12A cells, and for K14, SMA and caldesmon-1 only in MYO1089. The inhibition of GJIC by carbenoxolone resulted in a loss of cell-type specific staining. For all representative images of LCS, the white dashed lines delimit the position of the porous membrane, and MCF-12A are seeded on top and myoepithelial MYO1089 at the bottom of this membrane. A side view of the 3D representation of z-stacks of the whole membranes is shown in LCS and LCS + CBX. Nuclei were stained with DAPI. Scale bar = 10 μm.

Interestingly, when cultured together in LCS, both cell types showed improved differentiation based on marker expression. Luminal MCF-12A cells became negative for K14, α-SMA, and caldesmon-1, while remaining positive for K18 (Figure 5A, LCS). Similarly, myoepithelial MYO1089 cells lost expression of K18 but retained expression of the myoepithelial markers (Figure 5A, LCS). These results suggest that direct communication and interaction between the two layers promote more specific and differentiated phenotypes of these cell lines *in vitro*.

When direct communication was inhibited in the LCS using 100 µM CBX, cells lost this improved differentiation and co-expressed both luminal and myoepithelial markers, similar to the pattern observed in 2D monoculture (Figure 5A, LCS + CBX). Since Cx43 is the predominant connexin in both cell lines, these data suggest that Cx43 plays a crucial role in facilitating direct communication between the two principal cell types of the mammary epithelial bilayer. Inhibition of cytoplasmic exchange through Cx43 may result in impaired cell differentiation, leading to a luminal compartment that aberrantly expresses basal-like markers. While CBX did not completely altered RT-qPCR genes tested, it may suggest that GJIC is an important player for proper cell differentiation, but not the only one. Direct cellular contact also has a role in proper cell differentiation.

## DISCUSSION

There is a concerted global effort within the scientific community to accelerate discovery by reducing and replacing traditional animal studies with alternative models. These models aim to evaluate tissue differentiation mechanisms and assess the effects of chemicals and therapeutic compounds by focusing on molecular and cellular responses. Such approaches are increasingly used to predict toxicological outcomes relevant to human and environmental health. Among these innovations, complex co-culture systems and 3D culture models represent a significant advancement toward bridging the gap between conventional 2D cultures and *in vivo* studies.

In this study, we introduce a simple yet effective layered co-culture system that enables investigation of the dynamic interplay between luminal and myoepithelial cells in the mammary gland. Our findings demonstrate that direct communication via GJIC between MCF-12A and MYO1089 cells is critical for achieving appropriate cell differentiation and phenotypic specificity.

### Designing Complex *In Vitro* Models to Better Represent the Mammary Gland

Monolayer (2D) cell cultures have long served as the standard for research and preclinical testing, particularly in toxicology and pharmacology (Edmondson et al., 2014). However, numerous studies have shown that cells maintained under 2D conditions often fail to preserve the phenotypic characteristics of their tissue of origin (Nerger and Nelson, 2019). Consequently, they may yield misleading results in toxicological assessments and poorly predict *in vivo* responses (Knight and Przyborski, 2015).

In our study, cells cultured in the LCS, where direct communication between MYO1089 and MCF-12A is preserved, exhibited improved differentiation profiles compared to their 2D monoculture counterparts. These results suggest that incorporating such cell-cell communication is essential for modeling tissue-specific characteristics more accurately. These interactions profoundly influence cellular responses, gene expression, and the activation of various signaling pathways (Ravi et al., 2015).

Despite this, many existing 3D models of the mammary gland remain limited to monocultures of luminal cells or co-cultures with stromal cells (Lee et al., 2007, Koledova, 2017, Koledova et al., 2016, Hurtado et al., 2023, Marchese and Silva, 2012). Some *in vitro* bilayer epithelial models have been established using primary mouse or human mammary epithelial cells, or even whole-gland explants, yet these models pose challenges for mechanistic studies due to restricted opportunities for genetic manipulation (Florian et al., 2019). Furthermore, relatively few studies have employed myoepithelial cells in a 3D context, despite their critical role in secreting basement membrane components that support luminal cell polarization and differentiation (Weber-Ouellette et al., 2018, Sirka et al., 2018, Gudjonsson et al., 2005).

We propose that the LCS represents a fast, efficient, and reproducible platform where both luminal and myoepithelial cells maintain improved differentiation states. This model facilitates the study of luminal–myoepithelial interactions independently while preserving the reciprocal influence these cells types exert on each other, making it a valuable tool for studying mammary gland development, cancer progression, and response to environmental or therapeutic stimuli.

### Direct Communication Via Gap Junctions Supports Mammary Cell Differentiation

Cell-to-cell interactions mediated by gap, adherens, and tight junctions are critical for the normal development and function of the mammary gland (Gudjonsson et al., 2005, Sternlicht et al., 2006). We previously demonstrated that these junctions form a junctional nexus within the mammary gland epithelium, with a composition that changes across developmental stages, highlighting their fundamental role in regulating epithelial organization and function (Dianati et al., 2016). Correspondingly, studies from our group and others have shown that disruption of GJIC through a loss-of-function Cx43 mutation is associated with developmental abnormalities and functional impairments in the mammary gland (Plante et al., 2010, Plante and Laird, 2008). Despite this, only a limited number of *in vitro* models enable detailed investigation of the interactions between luminal and myoepithelial cells.

Using our LCS, we demonstrated that MCF-12A luminal cells and MYO1089 myoepithelial cells can establish direct intercellular contacts while residing on opposite sides of a porous membrane. This communication is likely mediated by cellular projections that traverse the pores. Within these membrane pores, we detected two key adherens junction proteins, E-cadherin and β-catenin, as well as functional gap junctions. The intracellular component of adherens junctions, β-catenin, binds to E-cadherins facilitating cell-cell adhesion and maintaining tissue integrity (Roura et al., 1999), playing an important role in mediating crosstalk between luminal and myoepithelial cells. β-catenin has a dual function as an anchorage protein and a transcription factor leading to the contribution of tissue morphogenesis and homeostasis (Tepera et al., 2003). It is involved in a variety of signaling processes and intercellular interactions, as thoroughly reviewed by Incassati et *al.* (Incassati et al., 2010).

E-cadherin is essential for maintaining the structural integrity of the bilayered mammary epithelium (Takekuni et al., 2003). It facilitates cell-cell adhesion between myoepithelial and luminal cells, forming a physical barrier that prevents cell intermingling and promotes proper spatial organization (Campbell et al., 2017). Although both luminal and myoepithelial cells express E-cadherin, its expression in myoepithelial cells is primarily localized to the ductal region rather than the lobular compartment (Hsiao et al., 2011), suggesting that our model most likely reflects the ductal architecture of the mammary gland. Moreover, general acini structures are composed of inner luminal cells surrounded by myoepithelial cells in a mesh-like structure (Gusterson et al., 1982), while, the ducts are structured as continued bi-layered. This reinforce that our model closely resembles the ductal anatomy of the mammary gland (Gusterson et al., 1982, Emerman and Vogl, 1986).

Additionally, we observed the presence of Cx43 within the membrane pores and confirmed functional communication between luminal and myoepithelial cells via gap junctions. Cx43 is the most ubiquitously expressed connexin in mammals and is predominantly localized to myoepithelial cells in the mammary epithelium, where it facilitates interactions with adjacent luminal cells (Dianati et al., 2016, Laird et al., 1999), a pattern recapitulated in our model. Beyond their canonical role in allowing passage of small molecules, connexins (including Cx43) interact with other proteins via their cytoplasmic tails (Dianati et al., 2017, Xu et al., 2001, Giepmans et al., 2001, Wei et al., 2005), enabling both GJIC-dependent and -independent mechanisms that regulate cell behavior and signaling in both cell types.

Animal model studies have shown that Cx43 is indispensable for proper mammary gland development and function (Plante et al., 2010, Plante and Laird, 2008, Stewart et al., 2013). In our LCS model, pharmacological inhibition of GIJC resulted in decreased levels of Cx43 and the loss of communication between MCF-12A and MYO1089 cells which in turn disrupted proper cell differentiation. Notably, Cx43 inhibition do not always affect the formation of tight or adherens junctions (Plante et al., 2010), indicating that gap junction-mediated cytoplasmic exchange via Cx43 is critical for maintaining epithelial cell polarization, differentiation, and function.

### Transcriptional Changes Induced by Direct Interaction Suggest Improved Cell Differentiation Compared to Classical 2D Cell Culture

Cell-cell communication significantly influences gene expression patterns, particularly when distinct cell types engage in direct contacts. To investigate how such interactions affect gene regulation, we performed RNA sequencing on luminal MCF-12A and myoepithelial MYO1089 cells. Interestingly, fewer genes were significantly altered in MYO1089 cells compared to MCF-12A, potentially due to differences in their baseline differentiation status. Indeed, while MCF-12A cells are considered non-tumorigenic breast luminal cells, they exhibit notable plasticity and do not express all typical luminal markers, thus being described as ‘pseudo-luminal’ cells (Sweeney et al., 2018). In contrast, MYO1089 cells display marker expression patterns consistent with primary myoepithelial cells (Holliday et al., 2009).

Notably, co-culture with MYO1089 led to the downregulation of genes associated with cell division and proliferation in MCF-12A cells. For example, SAPCD2, a protein involved in mitosis, showed decreased expression. SAPCD2 is typically expressed during embryogenesis but is markedly reduced in normal postnatal tissues (Baker and Du, 2022). The observed downregulation of genes involved in proliferation and the cell cycle suggests a transition away from a mesenchymal-like phenotype in MCF-12A cells, as active proliferation is generally incompatible with cellular differentiation (Brown et al., 2003, Ruijtenberg and van den Heuvel, 2016). Additionally, our results showed an upregulation of the Gene Ontology pathway associated with the regulation of animal organ morphogenesis, including genes implicated in cell polarization and differentiation. This further supports the hypothesis that myoepithelial cells contribute to luminal cell differentiation, likely through direct communication via Cx43.

In turn, MYO1089 cells in the co-culture condition displayed increased expression of K14, a key cytoskeletal protein essential for maintaining myoepithelial cytoarchitecture. K14 facilitates connections to neighboring cells through desmosomes and hemidesmosomes (Gudjonsson et al., 2002, Runswick et al., 2001). Desmosome-mediated adhesion is well known to regulate epithelial morphogenesis and cell positioning within tissues (Runswick et al., 2001, Qiu et al., 2008).

Furthermore, the main biological processes altered in myoepithelial cells during co-culture involved protein maturation and localization. Notably, we observed enrichment of pathways related to the regulation of vasculature development. In transgenic mouse models lacking vascular endothelial growth factor (VEGF), mammary glands displayed severely inhibited lobuloalveolar expansion during pregnancy and defective epithelial secretory activity during lactation (Qiu et al., 2008, Rossiter et al., 2007). This finding highlights the importance of vasculature regulation in supporting proper mammary gland development and function.

Taken together, our findings indicate that direct interactions between luminal and myoepithelial cells in the LCS model result in gene expression profiles more closely resembling those of a healthy mammary gland. The downregulation of basal-like and proliferation-associated genes, along with the upregulation of pathways related to proper tissue organization and differentiation, underscores the potential of this model to more accurately mimic luminal-myoepithelial dynamics *in vitro*.

It’s important to take into account that direct communication via gap junctions is important for proper cell differentiation but not the only factor. With our CBX experiment and RT-qPCR, we found that some genes followed classical 2D (in the LCS + CBX) but other closely resemble the healthy co-culture. While GJIC is blocked, direct physical interactions, may also help cellular differentiation. Indeed, physical contacts between adjacent cells can change signaling pathways and activation states (Bechtel et al., 2021). In our model, direct communication via gap junctions is crucial but not exclusive for proper cell differentiation.

### The Layered Co-Culture System: An Adaptable New Approach Methodology for Fundamental and Toxicological Research

This new model opens many perspectives for fundamental and toxicological research as cell-cell interactions play a central role in many tissues and biological processes. This methodology enables the *in vitro* evaluation of GJIC roles in proper cellular development, yielding results consistent with those observed in animal studies. As a modular and adaptable system, this model offers the ability to isolate and analyze individual components of the mammary gland’s complex architecture. Its versatility also allows for future incorporation of additional cell types, such as fibroblasts, adipocytes, cancerous cells, and extracellular matrix components, which will facilitate the investigation of the specific roles of each component of breast cancer. It is plausible that extending this model into a 3D format could further enhance its biological relevance. Indeed, 3D models are widely recognized for their ability to better replicate *in vivo* conditions, including cell proliferation, differentiation, and key interactions with the extracellular matrix (ECM). Moreover, the diverse set of analytical techniques used, including immunofluorescence on whole membranes or sections, RNA extraction, and dye-based functional assays, demonstrates the broad range of data that can be derived from this approach.

## CONCLUSION

We successfully developed a co-culture model that recapitulates key features of the mammary gland epithelium, demonstrating distinct more differentiated phenotypes for each cell type, and identifying junctional proteins formation and quantifying functional gap junctional activity between both layers. Overall, this co-culture model serves as a valuable platform for toxicological screening and mechanistic studies, supporting rapid risk assessment of agents that may affect the structural and functional integrity of both luminal and myoepithelial cells.

## Acknowledgements

We thank the Montreal Clinical Research Institute (IRCM) for their technical support and sample treatment. We thank Johanne Lorvinski for her technical assistance. We thank Michael G. Wade for his insightful feedback and review of the manuscript.

---

**Supplementary Figure 1.**
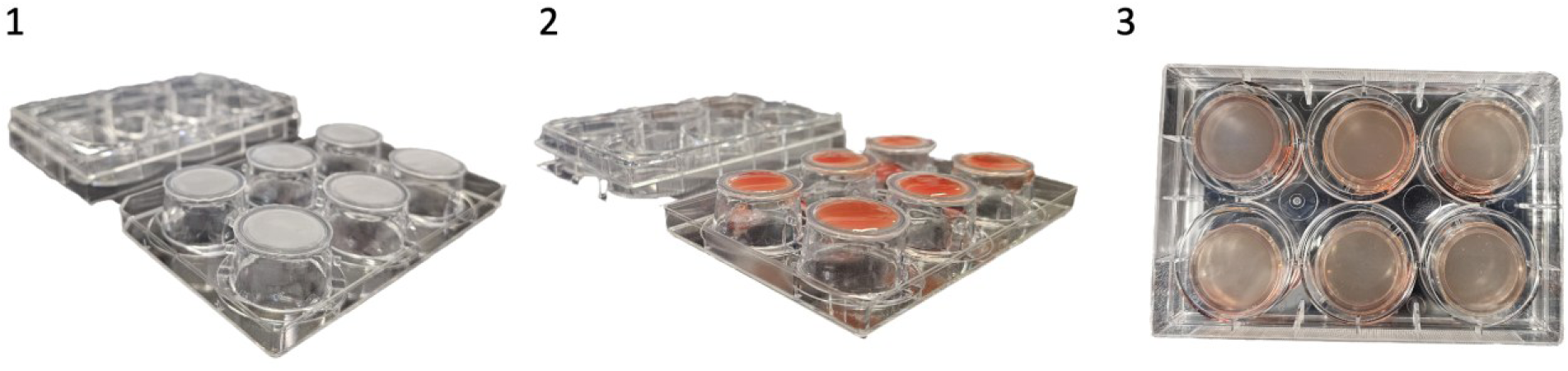
Images of the first steps of the layered-culture system protocol. (A) The procedure starts by placing the inserts up-side-down on the lid of a P6 plate. (B) Once the inserts are placed, 500 mL of MYO1089 cell suspension (1 million cells/mL) are placed in the center of the insert. (C) The P6 (wells) are used as a lid to hold in place by surface tension the droplet of MYO1089 cells. Then the system is transferred to an incubator at 37°C and 5% CO_2_ for six hours.

**Supplementary Figure 2.**
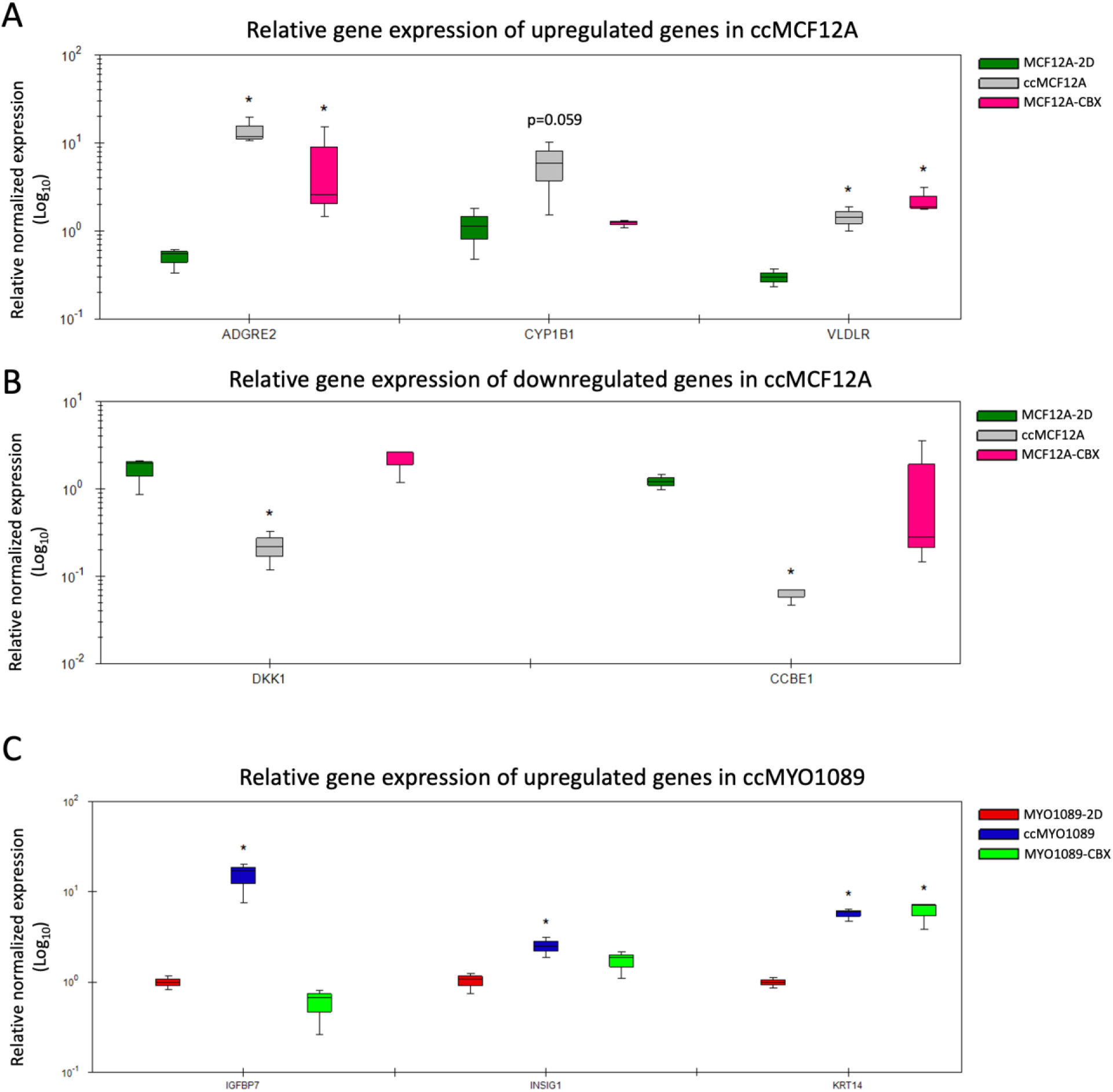
mRNA expression of genes affected by the co-cultured system and Carbenoxolone effect. (A) Illustrates the transcriptomic expression of upregulated genes ADGRE2, CYP1B1, and VLDLR, and (B) downregulated genes DKK1 and CCBE1 in the co-culture system exposed to 100 of μM carbenoxolone (CBX) or vehicle on MCF-12A cells. (C) illustrates the transcriptomic expression of upregulated genes IGFBP7, INSIG1, and KRT14 upregulated genes in the co-culture system exposed to of 100 μM carbenoxolone (CBX) or vehicle on MYO1089 cells. Graphs represent the mean +/-SD of three independent assays. Differences were considered significant when p≤0.05 following a T-test.

**Supplementary Figure 3.**
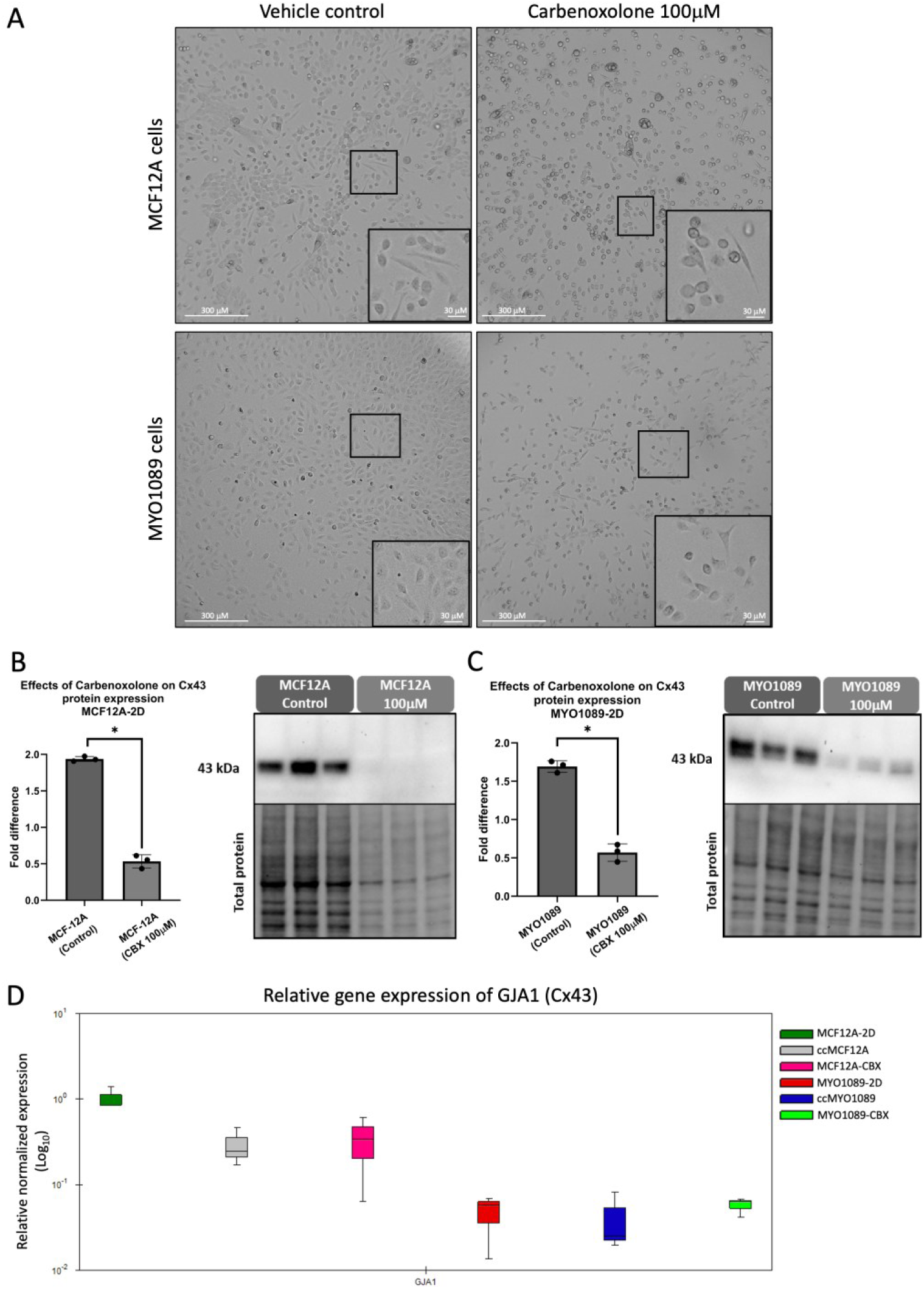
Carbenoxolone induces a shift in cell shape and a downregulation of Cx43 at protein levels. (A) When exposed to 100 μM carbenoxolone (CBX), MCF-12A cell shape predominantly shifts from an elongated phenotype to a rounder phenotype, leading to two distinct cell populations, without inducing cell toxicity. Semi-quantitative western blot analysis of total proteins extracted from 2D cell culture of MCF-12A (B) and MYO1089 (C) cells exposed to vehicle control or 100 μM of CBX. Histograms represent protein band normalized to the total protein level. Graphs represent mean +/-SD of Cx43 expression and total proteins used for loading normalization of three independent experiments. p-values were calculated with a Mann-Withney Test. Differences were considered significant when p≤0.05. (D) Illustrates the transcriptomic expression of GJA1 gene (Cx43) in the co-culture system and the effect of 100 μM carbenoxolone (CBX) in MCF-12A and MYO1089 cells.

**Supplementary Table 1.** Liste of Antibodies used for immunofluoresence

| Target protein | Host | Dilution | Manufacturer | Catalogue # |
| --- | --- | --- | --- | --- |
| **Primary antibodies** |  |  |  |  |
| β-catenin | Rabbit | 1:100 | Cell signaling | 8480 |
| E-cadherin | Rabbit | 1:100 | Cell signaling | 3195 |
| Cx43 | Rabbit | 1:250 | Sigma Aldrich | C6219 |
| K14 | Mouse | 1:100 | Epredia | ms-115-p1 |
| K18 | Rabbit | 1:100 | Abcam | ab52948 |
| αSMA | Rabbit | 1:100 | Abcam | 75635 |
| Caldesmon-1 | Rabbit | 1:100 | Cell signaling | 12503 |
| **Secondary antibodies** |  |  |  |  |
| Anti-mouse IgG<br>Alexa 488 | Goat | 1:1000 | Invitrogen | A11001 |
| Anti-rabbit IgG<br>Alexa 568 | Donkey | 1:1000 | Invitrogen | A10042 |

**Supplementary Table 2.**
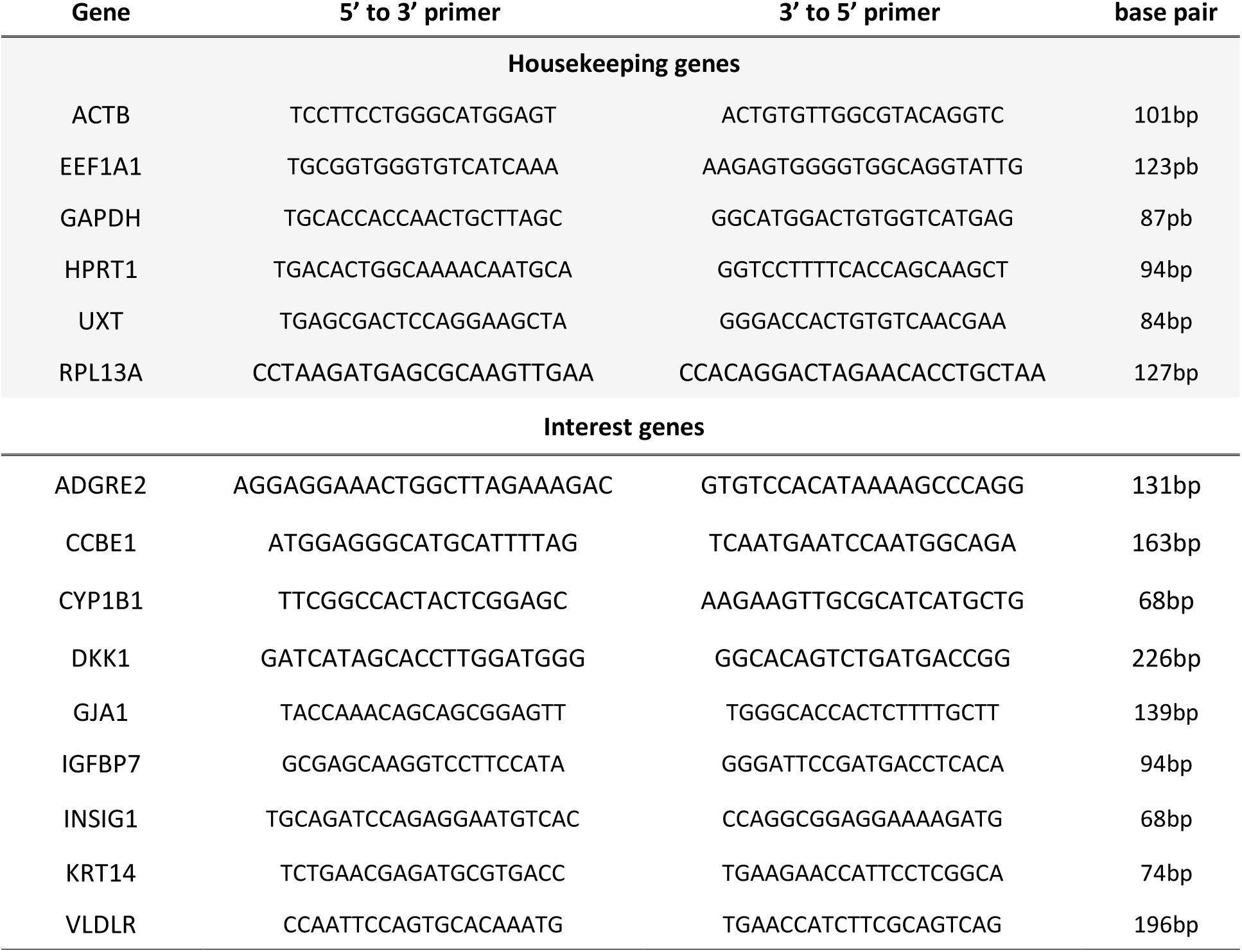
Liste of primers

